# Rational Design of a Bifunctional AND-Gate Ligand to Modulate Cell-Cell Interactions

**DOI:** 10.1101/711549

**Authors:** Jungmin Lee, Andyna Vernet, Katherine Redfield, Shulin Lu, Ionita C. Ghiran, Jeffrey C. Way, Pamela A. Silver

## Abstract

Protein “AND-gate” systems, in which a ligand acts only on cells with two different receptors, direct signaling activity to a particular cell type and avoid action on other cells. In a bifunctional AND-Gate protein, the molecular geometry of the protein domains is crucial. Here we constructed a tissue-targeted erythropoietin (EPO) that stimulates red blood cell (RBC) production without triggering thrombosis. EPO was directed to RBC precursors and mature RBCs by fusion to an anti-glycophorin A antibody V region. Many such constructs activated EPO receptors *in vitro* and stimulated RBC and not platelet production in mice but nonetheless enhanced thrombosis in mice and caused adhesion between RBCs and EPO receptor-bearing cells. Based on a protein-structural model of the RBC surface, we rationally designed an anti-glycophorin/EPO fusion that does not induce cell adhesion *in vitro* or enhance thrombosis *in vivo*. Thus, meso-scale geometry can inform design of synthetic-biological systems.

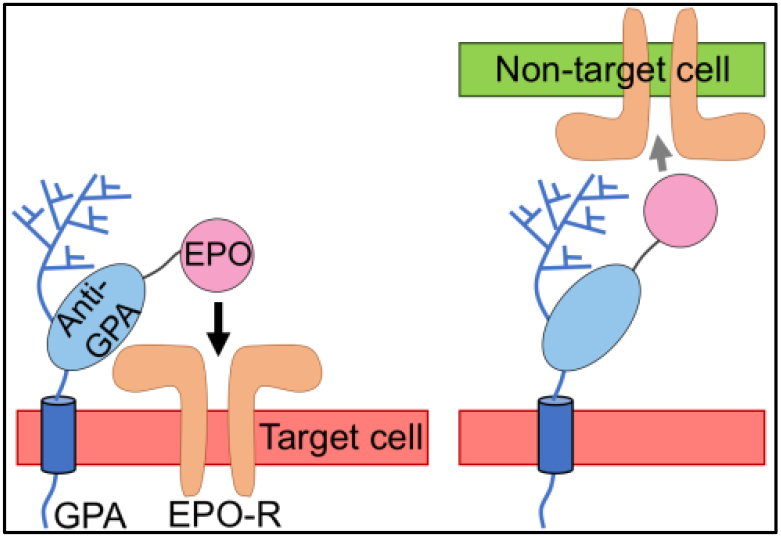

## INTRODUCTION

Hormones and cytokines act on receptors on diverse cells to achieve coordinated biological responses. Some activities are desired while others constitute drug side effects. For example, cortisol acts on the glucocorticoid receptor in immune cells and is anti-inflammatory, but acts on other cells to increase blood sugar and induce bone resorption. Similarly, erythropoietin (EPO) stimulates red blood cell (RBC) production but also increases thrombotic activity. These responses are adaptive in the context of hemorrhage, but increased thrombotic activity may underlie the increased frequency of heart attacks and strokes seen in EPO-treated kidney failure patients.

We previously designed a class of AND-Gate proteins, “Chimeric Activators,” in which a cytokine or hormone is mutated to reduce its activity, and is attached to a targeting element that binds to another protein on a subset of receptor-bearing cells (Figure 1a-d). Chimeric activators are similar to other fusion proteins such as blinatumomab*^1^* or immunocytokines,*^2^* which connect one cell to another. Thus, implicit in this design is the potential to crosslink cells, whether desired or not. Understanding the structural determinants for engineered cell-cell interaction will be important in design of next-generation CAR-T cells, bispecific T-cell engagers, and other fusion proteins.*^3^*

**Figure 1.**
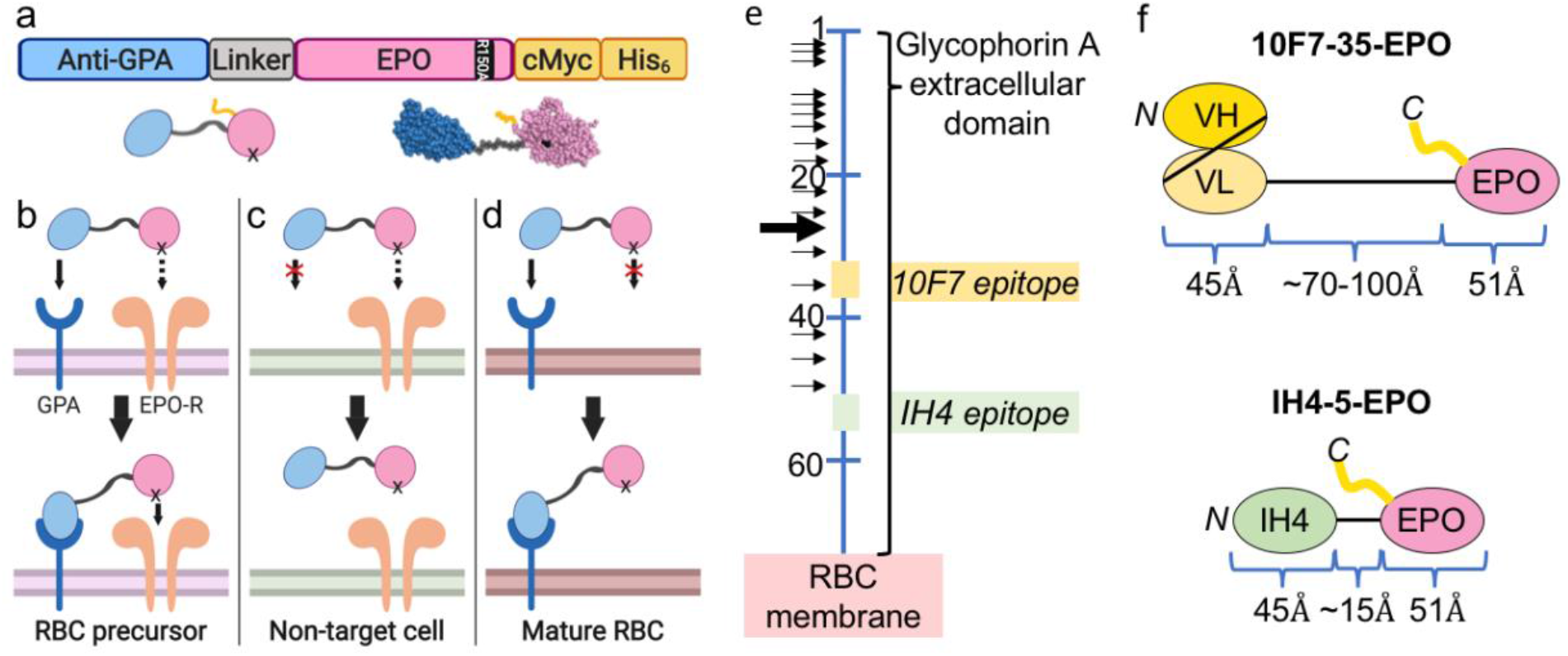
(a) Protein sequence organization of a Targeted EPO chimeric activator. (b) The chimeric activator binds to a target cell via the targeting element and tethers the mutated activity element to the cell surface, so that the mutated activity element binds to its receptor despite the mutation. (c) On a non-target cell, lacking the surface marker to which the targeting element binds, the mutated activity element has little effect. (d) Targeted EPO molecules also bind to mature RBCs via GPA, creating a sink for a Targeted EPO to extend its plasma half-life. EPO-R is not present on these cells. (e) The extracellular portion of GPA (blue line), showing amino acid positions (numbers), O-linked (small arrows) and N-linked (large arrow) glycosylation sites, and the positions of the 10F7 and IH4 epitopes. (f) The 10F7-35-EPO(R150A) and IH4-5-EPO(R150A) forms of Targeted EPO, showing sizes in Angstroms (Å) based on typical scFv and nanobody sizes, an extended conformation of the Gly/Ser linkers, and solved structures of EPO. Some images were created with BioRender.

We previously constructed an AND-gated EPO that activates EPO receptors (EPO-Rs) on RBC precursors but not on other cells.*^4,5^* “Targeted EPO” proteins consist of an antibody element that binds to glycophorin A (GPA), EPO with a mutation that reduces EPO activity, and a connecting linker (Figure 1a).^*4,5*^ GPA is abundant on RBCs (800,000 copies/cell) and late RBC precursors (50,000 copies/cell).*^4,6^* GPA is not expressed on cells that control thrombosis and tumor angiogenesis, or solid tumor cells. EPO-Rs on late RBC precursors mediate EPO-dependent maturation, but then disappear.*^4,7^* Targeted EPO was designed to bind to late RBC precursors and stimulate EPO-Rs on only those cells (Figure 1b). On non-target cells, it fails to activate EPO-Rs because the mutated EPO alone binds poorly to receptors (Figure 1c). Binding to GPA on mature RBCs is a desirable secondary consequence as it extends the plasma half-life of the fusion protein.*^4^* However, EPO bound to mature RBCs could interact with EPO-Rs on endothelial cells*^8^* or immune cells such as macrophages*^9^* and mediate undesired cell-cell adhesion or signaling (Figure 1d).*^4^*

## RESULTS AND DISCUSSION

The present work defines structural features of anti-GPA/EPO chimeric activators designed to act specifically *in cis* on GPA-expressing cells, to avoid side effects that may arise from unwanted cell-cell interaction. Previously we characterized a form of Targeted EPO consisting of V regions from the 10F7 antibody, a 35-amino acid linker, and EPO with the weakening mutation R150A; this version is termed 10F7-35-EPO(R150A) (Figure 1f). This protein specifically stimulated production of RBCs but not platelets, the latter being considered an indicator of off-target cell effects. To directly assess the impact of this molecule on thrombosis, we performed a ‘bleeding time’ assay in which the tail of a treated mouse is transected and the time to cessation of bleeding is measured.

Contrary to expectation, the 10F7-35-EPO(R150A) form of Targeted EPO promoted thrombosis in the bleeding time assay (Figure S1). We hypothesized two mechanisms by which this might occur. First, the antibody element could, by itself, promote blood clotting. Second, the RBC-bound Targeted EPO could crosslink RBCs with other cells bearing EPO-Rs, such as vascular endothelial cells or leukocytes, slowing blood flow. To determine whether either or both of these hypotheses were correct, we constructed a series of Targeted EPO variants, testing anti-GPA antibody elements (10F7, 1C3, R18 and IH4), fusion protein configurations, and linker lengths.

### Targeted EPO activates EPO-Rs in a GPA-dependent manner in vitro and in vivo

Diverse forms of Targeted EPO activate EPO-Rs in a GPA-dependent manner *in vitro*. TF-1 cells, which express both GPA and EPO-Rs,*^5^* were treated with several forms of Targeted EPO, and their proliferation was measured 72 hr post-treatment (Figure 2a,b). Unfused EPO(R150A) showed about 100-fold reduced activity compared to wild-type EPO (EPO(WT)). When EPO(R150A) was fused to an anti-GPA antibody element, its activity was rescued by about 10-100-fold (Figure 2a,b). IH4 and 1C3 were the most potent antibody elements in these tests.

**Figure 2.**
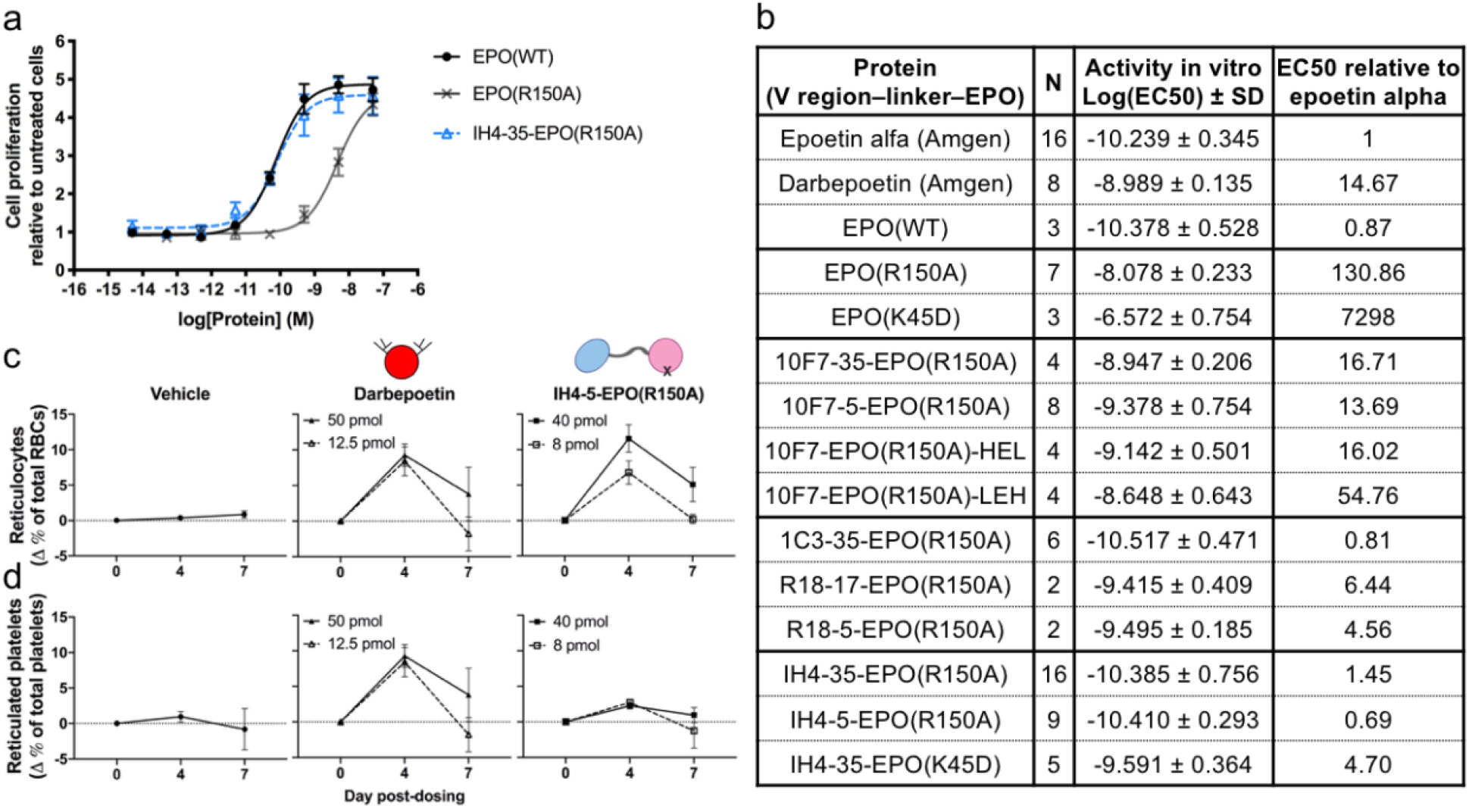
*In vitro* and *in vivo* activity of different forms of Targeted EPO. (a) A typical TF-1 cell proliferation assay comparing the stimulation of proliferation by “wild-type” EPO (including C-terminal tags), the EPO(R150A) mutant protein, and the IH4-35-EPO(R150A) fusion protein. The effective concentration (EC50) is calculated by a 4-parameter fit. (b) A table summarizing the ability of different forms of Targeted EPO and control proteins to promote proliferation of TF-1 cells *in vitro*. (c,d) An *in vivo* test of the ability of IH4-5-EPO(R150A) to specifically promote RBC production and not platelet production, using transgenic mice that express human GPA on their RBCs.*^4^* The results indicate that IH4-5-EPO(R150A) and diverse other forms of Targeted EPO show enhanced activity of the EPO(R150A) element independently of the mode of attachment of EPO to the GPA-binding antibody element. Data represent mean ± SEM.

Three configurations of Targeted EPO were tested to address whether restricting the movement of EPO would affect EPO activity and potentially cause GPA-bound EPO to preferentially act *in cis* on targeted cells. The conventional scFv-EPO configuration is designed as antibody heavy chain – light chain – EPO (“HLE”). We modified it such that EPO would be between the heavy and light chains; this is feasible because the N- and C-termini of EPO are close in the folded structure. The resulting configurations (“HEL” and “LEH”) were expressed and tested. Both 10F7-EPO(R150A) and 1C3-EPO(R150A) stimulated proliferation of TF-1 cells largely independently of the configuration, indicating that orienting EPO in one way does not significantly constrain its receptor binding *in cis* (Figure 2b and Table S1).

The GPA-mediated enhancement of EPO(R150A) activity was independent of the linker length in the various fusion proteins. Changing the linker of 10F7-EPO(R150A), R18-EPO(R150A), R18-EPO(K45D), IH4-EPO(R150A), and IH4-EPO(K45D) from 5 to 35 amino acids (about 10-80 Angstroms) in fusion proteins containing the same antibody and EPO elements showed similar EC50s in cell-based assays (Figure 2b and Table S1). Results with the rotationally constrained and short-linker constructs are consistent with the idea that GPA is an intrinsically disordered, highly flexible protein, so that diverse fusion proteins bound to GPA can still allow binding of EPO with EPO-R.

When tested *in vivo*, various forms of Targeted EPO specifically stimulated production of RBCs but not platelets. We assessed *in vivo* target cell specificity and efficacy of several Targeted EPO proteins in mice that are transgenic for human GPA (following Burrill *et al*.*^4^*). Darbepoetin, an extended half-life form of EPO, is a control to represent non-targeted EPO activity. Mice were injected intraperitoneally with vehicle, darbepoetin, or a Targeted EPO, and their reticulocytes and reticulated platelets were measured to indicate activity on target cells and non-target cells, respectively. Mice treated with darbepoetin showed comparable increases in both new RBCs and new platelets (Figure 2c,d and S2).*^4^* Mice treated with IH4-5-EPO(R150A) showed a similar increase in reticulocytes while the reticulated platelets were not significantly changed (Figure 2c,d). IH4-5-EPO(R150A) was the most potent *in vivo* of the forms of Targeted EPO tested (Figure 2b and S2).*^4^*

### The antibody element IH4 does not induce ‘RBC inflammation’

The anti-GPA antibody element in forms of Targeted EPO might induce an inflammatory phenotype that could enhance blood clotting or cause other undesired side effects and also confound interpretation of experimental treatment with fusion proteins. Various anti-GPA antibodies and V region elements induce a constellation of effects on RBCs, including increased stiffness during shear stress, decreased membrane fluidity, secretion of ATP, production of reactive oxygen species, and phosphorylation of Tyr8 of band 3. Divalent IgG-type antibodies against GPA induce at least some and likely all of these phenotypes, but only a subset of monovalent Fabs, scFvs and VHHs induce this RBC response in an epitope-dependent manner (Figure 1e).*^10–15^*

The single-chain camelid element (a VHH or nanobody) IH4 itself lacks stiffening/inflammatory activity when it is bound to GPA, and therefore is a good candidate targeting element.*^13^* This element binds to the membrane-proximal epitope (_52_YPPE_55_) on human GPA.*^14^* We tested an optimized variant of IH4 (Figure S3). We confirmed by fluorescence recovery after photobleaching (FRAP) that 10F7 induced lower membrane lipid mobility in RBCs, while IH4 did not (Figure 3). In addition, IH4 appears to bind most tightly of the known anti-GPA V region elements. Therefore, IH4 was chosen as a candidate antibody element for a Targeted EPO that would not induce undesired signaling and would also limit potential interaction of EPO with EPO-Rs on other cells.

**Figure 3.**
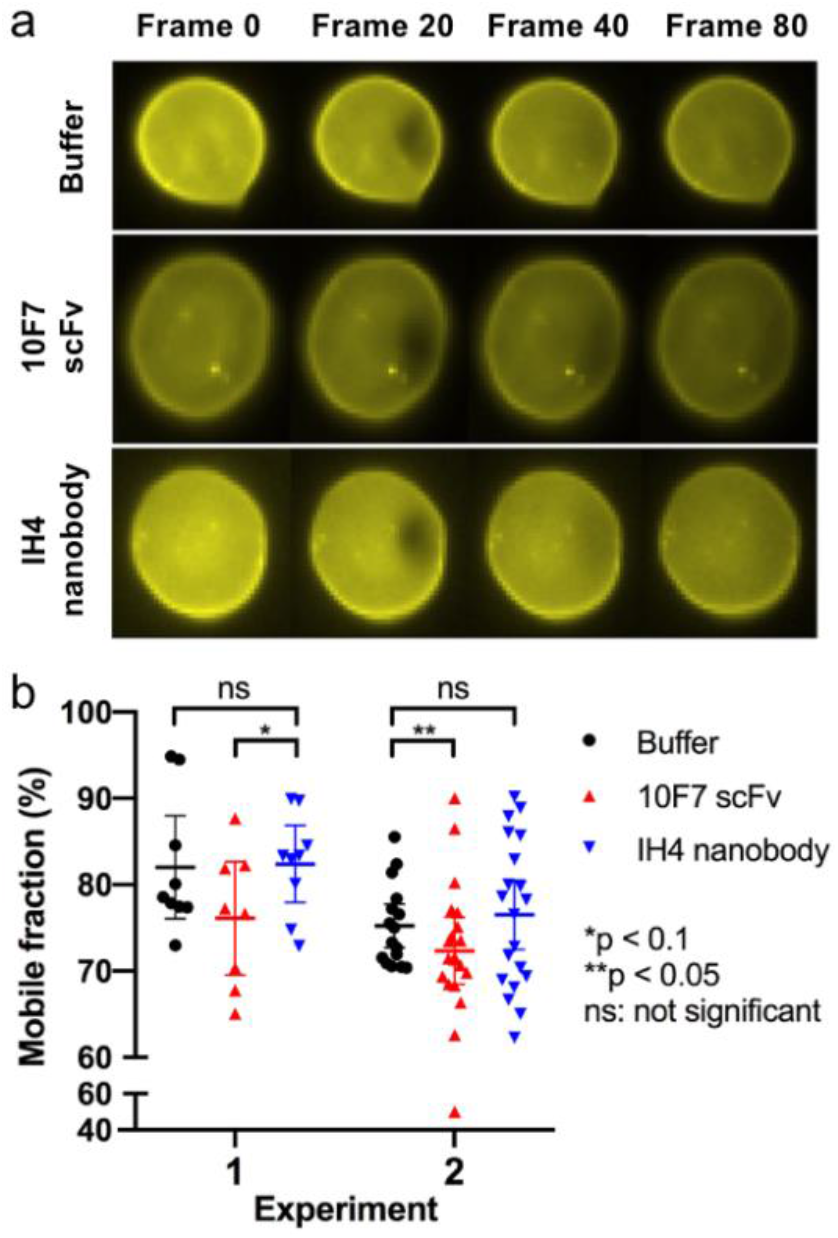
Reduction in membrane fluidity by 10F7 but not IH4, measured by FRAP. RBC membranes are stained with a lipid-soluble dye, a portion of the membrane is bleached with a laser, and diffusion of the dye into the bleached region is measured. (a) Serial snapshots of RBCs treated with antibody fragments during FRAP experiments. Each recording was made at a rate of 8 frames per second for 10 seconds. (b) Mobile lipid fraction of 8-9 RBCs (Experiment 1) or 16-20 RBCs (Experiment 2) treated with buffer, 10F7, or IH4. When the data from these experiments were normalized to the buffer controls in the same experiments and then combined, the difference between 10F7 and buffer (p = 0.0648) or 10F7 and IH4 (p = 0.0463) was significant (Mann-Whitney test). Lines indicate mean and 95% confidence interval. The distribution of individual cell responses is typical for these assays.*^16^*

### Geometry of GPA and Targeted EPO determines potential for cell crosslinking

In the design of Targeted EPO, the intention is that the antibody element and EPO bind *in cis* to receptors on the same cell surface (Figure 1b). However, fusion proteins comprising two binding elements have the potential to crosslink different cells that each bears a receptor for a component of the fusion protein. To address this, we tested the accessibility of antibodies to the EPO element of Targeted EPO variants bound to RBCs, and the ability of different Targeted EPOs to promote binding between RBCs and tumor cells expressing EPO-Rs.

When forms of Targeted EPO are bound to RBCs via GPA, the antibody accessibility of the EPO element decreases when the linker length is shortened and the epitope on GPA is closer to the cell membrane. We compared a series of proteins in which EPO was fused to IH4 or 10F7 and the linker length was varied. The proteins had a His_6_ tag at the C-terminus. RBCs were incubated with the fusion protein, and then the accessibility of EPO in the bound state was assessed via flow cytometry by binding of a phycoerythrin (PE)-conjugated anti-His_6_ antibody. For both IH4 and 10F7 fusion proteins, PE signals progressively decreased as the linker was shortened, suggesting that the fusion proteins with shorter linkers were better masked from interacting with the antibody (Figure 4a). In addition, crosslinking of RBCs by the divalent anti-His_6_ antibody increased with the linker length of the fusion protein, as seen in forward scattering signals evidenced by a shoulder to the right of the main peak (Figure 4b). Also, regardless of the linker length, all IH4 fusion proteins had lower accessibility and fewer RBC crosslinking events by anti-His6 antibody than corresponding 10F7 fusion proteins, suggesting that the binding epitope of the antibody element also affects the accessibility of EPO (Figure 4a,b).

**Figure 4.**
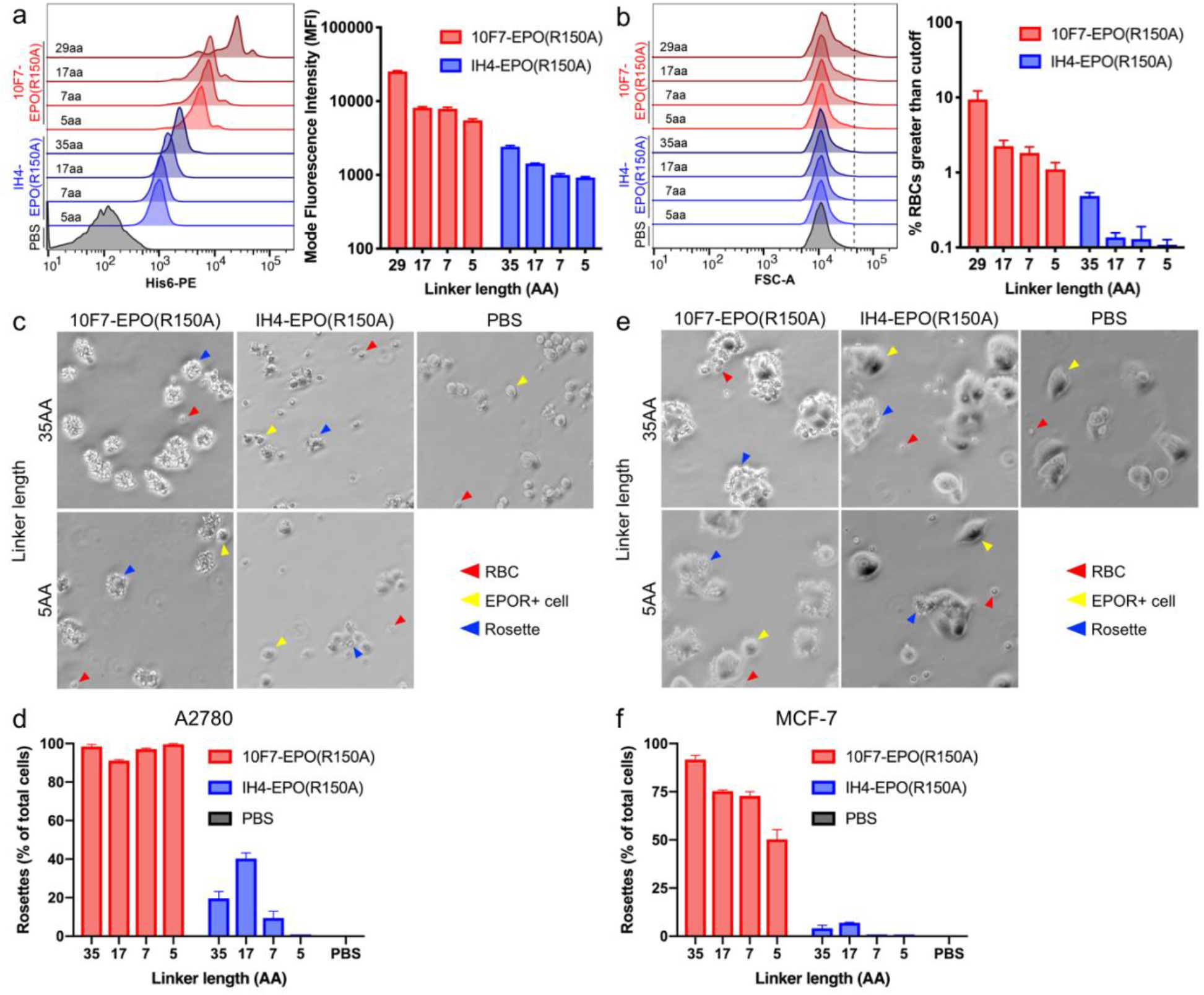
Targeted EPO fusion proteins with short linkers and membrane-proximal GPA binding epitopes reduce EPO accessibility and cell crosslinking. (a,b) Forms of Targeted EPO containing either 10F7 or IH4 and varying linker lengths were incubated with human RBCs, and stained with a PE-labeled anti-His_6_ antibody. (a) The accessibility of the EPO element by anti-His_6_-PE antibody is shown as mode fluorescence intensity (PE). (b) The crosslinking of RBCs mediated by divalent anti-His_6_ antibody is shown as forward scattering (particle size). The signal greater than the cutoff (dotted line) indicates adhesion of RBCs. (c-f) Human RBCs treated with Targeted EPO variants were incubated with (c,d) A2780 or (e,f) MCF-7 cells, both of which express EPO-R. (c,e) Phase-contrast images of RBC rosetting around EPO-R-bearing cells. 200x magnification. (d,f) The frequency of rosette formation was scored as tumor cells bound by 3 or more RBCs, shown as a percentage of total tumor cells. Data represent mean ± SEM.

Forms of Targeted EPO with long linkers and a membrane-distal epitope on GPA mediate adhesion *in vitro* between RBCs and cells expressing EPO-Rs. We used a rosette assay for cell-cell interaction to address whether the anti-GPA/EPO fusion proteins might promote adherence of RBCs to cells from the tumor lines A2780 and MCF-7, which express EPO-R.*^17,18^* All forms of Targeted EPO containing the 10F7 scFv (epitope at about _34_YAATP_38_) showed a high level of rosette formation, with most or all of the tumor cells binding to 3 or more RBCs. In contrast, the frequency of rosette formation with Targeted EPOs containing the IH4 nanobody (epitope: _52_YPPE_55_) was much lower. Targeted EPO forms with shorter linkers generally induced less rosette formation. These effects were synergistic: the IH4-EPO(R150A) with a 5 amino acid linker showed essentially no rosette formation by either A2780 or MCF-7 cells (Figure 4c-f and S4).

The results on antibody accessibility and rosette formation are consistent with a geometric model of the RBC surface in which GPA extends from the cell and shields the cell from interactions with other cells. GPA is densely packed on the RBC surface, with about 800,000 monomers*^6^* in a surface area of 140 μm^2^,*^19^* such that the average distance between GPA molecules is about 130-190 Angstroms (Figure S5). The membrane-distal N-terminal ~45 amino acids of GPA include 17 sites for O- and N-glycosylation, with oligosaccharides that extend at least about 20 Angstroms from the peptide backbone (for the O-type blood group antigen). It is thought that the N-terminal 45 amino acids of GPA are largely unstructured and would be about 90-130 Angstroms long in an extended conformation.*^20^* By comparison, in an IgG antibody in the T conformation the tips of the V regions are about 150 Angstroms apart, and in an scFv-(SGGGS)-EPO fusion protein, the components would be 45, 15 and 51 Angstroms end-to-end. Thus, antibody access to membrane-proximal epitopes could be limited, and contact between a GPA-bound IH4-(SGGGS)-EPO molecule and EPO-Rs on another cell should be sterically very difficult (Figure 1e,f).

### IH4-5-EPO(R150A) does not shorten tail vein bleeding time in mice

We tested IH4-5-EPO(R150A) for systemic effects on thrombosis in mice. HuGPA-transgenic mice were injected ip with vehicle, darbepoetin, 10F7-35-EPO(R150A) or IH4-5-EPO(R150A) (Figure 5a). EPO proteins were injected at a dose that resulted in similar effect in mice, as measured by reticulocyte counts. In mice treated with darbepoetin or 10F7-35-EPO(R150A), the bleeding time was shortened by about 30%, whereas mice treated with IH4-5-EPO(R150A) showed no change in bleeding time (Figure 5b,c and S1). These results indicate that IH4-5-EPO(R150A) does not enhance thrombotic side effects, in contrast to the non-targeted form EPO that acts on EPO-Rs on multiple cell types. When the data from these experiments were normalized to the vehicle controls of the same experiments and then combined, the difference between vehicle and darbepoetin (p = 0.0098) or IH4-5-EPO(R150A) and darbepoetin (p = 0.017) was significant. When data were compared across all experiments with dose groups including 10F7-35-EPO(R150A) and epoetin alfa, these molecules also showed a statistically significant decrease in bleeding time relative to the vehicle control (Figure S1). These results are consistent with previous studies indicating that non-targeted EPO proteins reduce bleeding time in rodents.*^21^*

**Figure 5.**
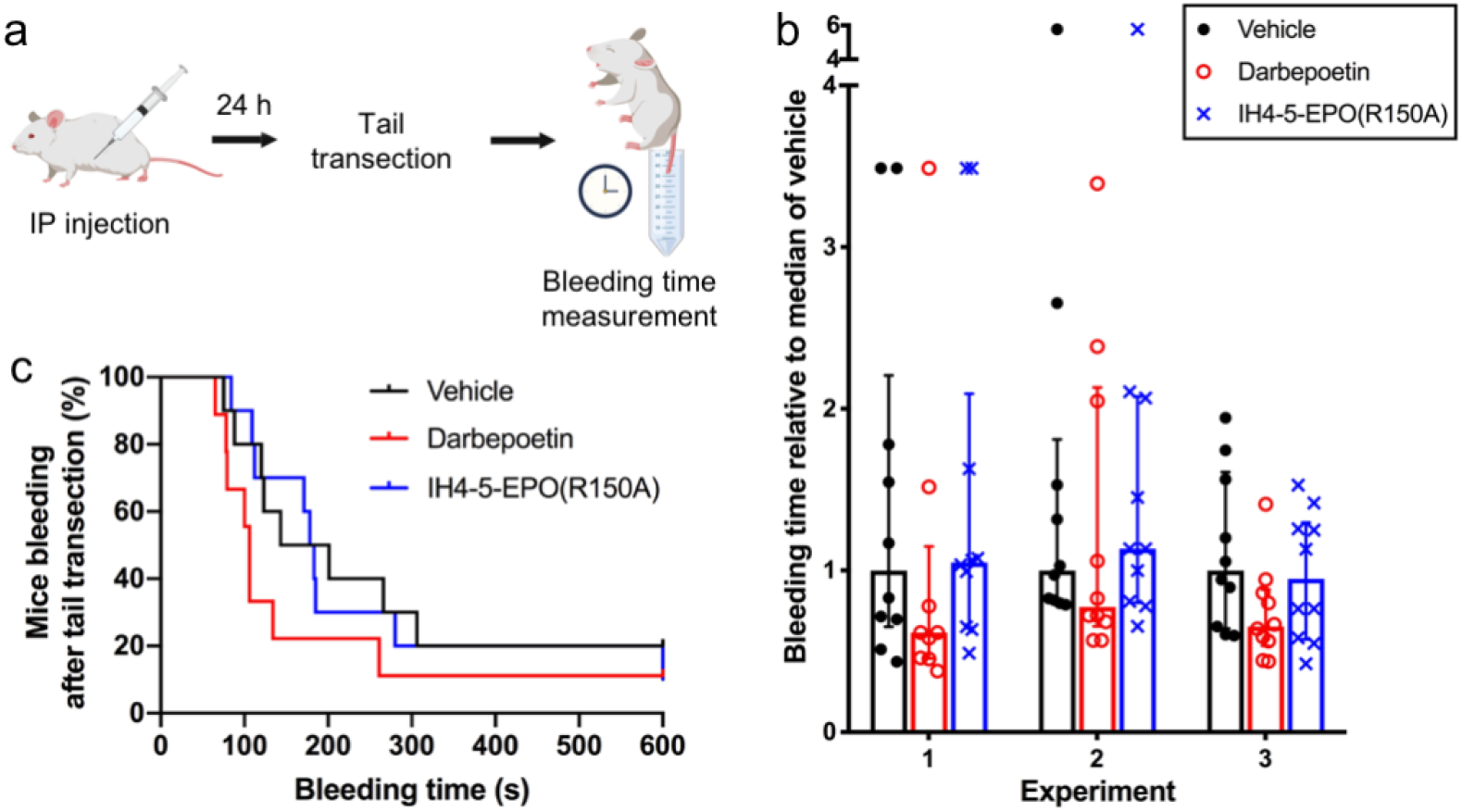
Shortened tail bleeding time in mice treated with a non-targeted EPO but not with IH4-5-EPO(R150A). (a) Experimental approach. Mice received a single ip injection of darbepoetin (50 pmol; 1.8 μg), IH4-5-EPO(R150A) (40 pmol; 2 μg), or vehicle. 24 hours later, tail transection was performed and bleeding times were recorded (See Methods). (b) Three experiments showing bleeding times relative to the vehicle control in huGPA-transgenic mice treated with vehicle, darbepoetin, or IH4-5-EPO(R150A). Data represent median with interquartile range. (c) A Kaplan-Meier plot for Experiment 1 from the data shown in (b).

### Conclusions

The design of a fusion protein must consider the spatial orientation of the target proteins to function while minimizing unwanted interactions. Such considerations become important in AND-Gate bifunctional protein systems that bind to different receptors abundant on different cell types. EPO-Rs are found on numerous cell types besides RBC precursors, including platelet precursors, vascular endothelial cells, liver cells, and white blood cells such as macrophages.

Here we addressed this problem for “Targeted EPO” molecules, fusion proteins that bind to GPA and weakly to EPO-R and are thus AND-gated to bind to only cells with both of these transmembrane proteins – namely late RBC precursors. The design goal is to specifically activate EPO-R only on late RBC precursors as a result of initial binding to GPA, followed by binding *in cis* to EPO-R due to the high local concentration of EPO (Figure 1b). We found that:

1. Some Targeted EPO proteins mediate adhesion between mature RBCs that express only GPA and other cells that express only EPO-R.
2. The position of the epitope on GPA and the length of the linker between the GPA binding element and the EPO element determines whether a given Targeted EPO can mediate cell adhesion.
3. The cell adhesion results can be interpreted with a structural model in which GPA extends from the RBC surface and limits how close another cell can approach. To mediate cell adhesion, the EPO element must be able to extend beyond the membrane-distal end of GPA.
4. More broadly, cell-cell interaction may be regulated by abundant cell surface proteins that may sterically prevent close approach of cells. This needs to be considered when constructing engineered cell interaction systems, such as CAR-T cells and bispecific antibodies.

A major challenge for synthetic biology is that artificial biological systems have the potential for undesired interactions that can lead to system failure. When we study natural systems, we are generally not aware of all the behaviors they could have if they were poorly designed. Our experiments illustrate geometric principles that natural systems may use in achieving specificity.

## METHODS

Methods for cell culture, DNA construction, protein expression and purification, *in vitro* measurement of erythropoietin activity by cell proliferation assays, and measurement of mouse reticulocytes and reticulated platelets are described in Burrill *et al.^4^* Fluorescence recovery after photobleaching (FRAP) was performed as described by Khoory *et al.^16^* See Supplementary Methods.

### EPO accessibility and RBC crosslinking measurement by flow cytometry

RBCs (Zen-Bio, Research Triangle Park, NC; 0.2% v/v in PBS-glucose) were incubated with fusion proteins for 1 hr at 4°C, washed once with PBS-glucose (PBS, 5 mM glucose, pH 7.4), incubated with anti-His6-PE antibody (Abcam) for 30 min at 4°C, and washed once with PBS-glucose. Fusion protein concentration used was calculated based on the K_D_ of the monovalent 10F7 and IH4 antibody elements (95 nM and 33 nM, respectively) to achieve 80% saturation of GPA. The cells were analyzed by standard flow cytometry techniques (see Supplementary Methods).

### RBC rosette assay

A2780 and MCF-7 cells were plated on a 6-well plate containing 12 x 12 mm coverslips and incubated overnight at 37°C. Next day, a mixture of RBCs (0.2% v/v) and fusion proteins was prepared in RPMI-1640 without bicarbonate and incubated for 30 min at 23°C. Coverslips with cells were washed twice with PBS and transferred to a new 6-well plate. The RBC-fusion protein mixture was added on top of each coverslip and incubated for 1 hr at 37°C. Coverslips were gently washed twice with PBS. Rosettes were imaged on a Nikon Eclipse TE300 inverted phase-contrast microscope. Rosettes were defined as adherent cells bound by 3 or more RBCs among at least 160 cells from at least 8 fields. See Supplementary Methods for further details.

### Mouse tail vein bleeding time measurement

All mouse experiments were performed under the protocol IS00000723, approved by the Harvard Medical School IACUC. Experiments were performed in a blinded manner. Ten mice per dose group received a single ip injection with saline or test protein. Next day, mice were anaesthetized. The tail was transected 3 mm from the tip, and the body of the mouse was placed on a heated pad over a 37 °C water bath containing tubes with 50 mL of saline, such that 0.5-2 mm of the transected tail was placed in the saline within 2 seconds of transection. The bleeding time was recorded when bleeding stopped for one minute. Bleeding time longer than 10 minutes were recorded as 10 minutes. See Supplementary Methods for detailed description.

## ACKNOWLEDGMENTS

This work was supported by funds from the Wyss Institute for Biologically Inspired Engineering and the Boston Biomedical Innovation Center (Drive Award).

