## Supplementary material for "Rational Design of a Bifunctional AND-Gate Ligand to Modulate Cell-Cell Interactions": Supplemetnary Information

### SUPPORTING INFORMATION

Jungmin Lee,[*,1,2] Andyna Vernet,[2] Katherine Redfield,[3] Shulin Lu,[4] Ionita C. Ghiran,[4] Jeffrey C. Way,[*,2,6] and Pamela A. Silver[2,5,6]

[1]Department of Chemistry and Chemical Biology, Harvard University, Cambridge, MA 02138, United States
[2]Wyss Institute for Biologically Inspired Engineering, Harvard University, Boston, MA 02115, United States
[3]Harvard-MIT Division of Health Sciences and Technology, Cambridge, MA 02139, United States
[4]Department of Medicine, Beth Israel Deaconess Medical Center, Harvard Medical School, Boston, MA 02115, United States
[5]Department of Systems Biology, Harvard Medical School, Boston, MA 02115, United States
[6]These authors contributed equally to this work.
*Corresponding authors

### Supplementary Methods

#### Cell culture

FreeStyle 293-F, FreeStyle CHO-S, and CHO DG44 cells were obtained from Invitrogen (Carlsbad, CA), and were cultured in FreeStyle 293 Expression Medium, complete FreeStyle CHO Expression Medium, and complete CD DG44 (Invitrogen), respectively. Human erythroleukemia TF-1, human ovarian cancer A2780, and human breast cancer MCF-7 were obtained by ATCC (Manassas, VA). TF-1 was cultured in RPMI-1640 with 10% FBS, 100 U/mL penicillin, 100 U/mL streptomycin, and 2 ng/mL recombinant human granulocyte macrophage colony-stimulating factor (GM-CSF; PeproTech) unless specified otherwise. A2780 was cultured in RPMI-1640 with 10% FBS. MCF-7 was cultured in DMEM with 10% FBS and 0.01 mg/mL recombinant human insulin (PeproTech). 293-F, CHO-S, and CHO DG44 were cultured at 37 °C in 8% $CO_2$ with shaking at 2.35 × g. TF-1, A2780, and MCF-7 were cultured at 37 °C in 5% $CO_2$.

#### DNA constructs

The DNA sequences for 10F7 and EPO (wildtype and mutants) were from GenBank (accession no. KX026660-3). The DNA sequences for IH4, R18, and 1C3 were derived by reverse translating and codon optimizing (Integrated DNA Technologies) the protein sequences adapted from the US patent 9879090, US patent 8900592, and patent application WO1993024630, respectively.[1-3] The IH4 sequence was modified to include a point mutation (Phe80Tyr) and an additional amino acid (Thr118) in the framework regions 3 and 4, respectively, as the reported sequence of IH4 may have had fortuitous changes from the germline sequence or typographical errors. The IH4 antibody with the original sequence in the patent was denoted as IH4*. The DNA sequences for glycine-serine linkers of various lengths were codon-optimized for expression in mammalian cells.

#### Protein expression and purification

Transient expression was performed in 293-F and CHO-S cells using pSecTag2A or pOptiVEC plasmids, and stable expression was performed using CHO DG44 cells using pOptiVEC plasmids according to the supplier's protocol. Protein expression was assayed by Western blotting cell supernatant using anti-$His_6$-HRP antibody (Abcam). Stably transfected cells were selected by hypoxanthine/thymidine (HT)-deficient CD OptiCHO medium (Invitrogen), and were subjected to one round of methotrexate (MTX; Sigma-Aldrich) genomic amplification as described before.[4] Proteins from transient transfection or stable pools were purified as follows. Supernatant was concentrated to 5-8 mL using a 10 kDa cut-off Macrosep Advance centrifugal device (Pall). Concentrated protein was bound to 0.5-1 mL of His60 nickel or HisTalon cobalt resin (Takara Bio) for 0.5 h at 4°C while rotating in a 10-mL Pierce disposable column (Thermo Scientific), and was washed and eluted using His60 or HisTalon Buffer Set (Takara Bio) according to the supplier's protocol. Cell supernatant and each purification fraction were analyzed by SDS-PAGE followed by Coomassie Blue staining. Eluted proteins were combined, desalted into endotoxin-free PBS (Teknova: 137 mM NaCl, 1.4 mM $KH_2PO_4$, 4.3 mM $Na_2HPO_4$, and 2.7 mM KCl, pH 7.4) using Econo-Pac 10DG columns (Bio-Rad), and concentrated to <1 mL using Macrosep Advance centrifugal device. For *in vivo* experiments, contaminating proteins were further removed by size exclusion chromatography (SEC) on Superdex 200 10/300 GL columns (GE Healthcare) using AKTA FPLC system (GE Healthcare) and endotoxin-free PBS as the running buffer. Desired protein fractions were combined and concentrated to <1 mL using Macrosep Advance centrifugal device. Proteins were stored at 4°C throughout the described process, ultimately stored as aliquots at −80°C, and thawed once before use. Only endotoxin-free reagents were used.

#### Cell proliferation assays

TF-1 cells were seeded in a 96-well plate at $9.0 \times 10^3$ cells per well in 90 μL of RPMI-1640 with serum and antibiotics (no GM-CSF). The purified proteins were serially diluted by 10-fold ($10^{-7}$ to $10^{-14}$ M) and added to the cells. After 72 hours, cell proliferation was measured by CellTiter 96® AQueous One Solution Cell Proliferation Assay (Promega). Absorbance at 490 nm was read on a BioTek Synergy Neo HTS microplate reader. Reported data represent mean ± SEM of three replicates.

**Flow cytometry for reticulocyte and reticulated platelet counts**

HuGPA transgenic FVB mice were generously donated by the Hendrickson Laboratory at Emory University.[5] This strain underwent embryo re-derivation at Charles River Laboratories. The homozygous huGPA transgene is embryonic-lethal but heterozygotes are phenotypically normal, so a breeding colony was maintained with screening for huGPA at each generation. Transgene expression was measured as described before.[4]

In experiments that measure efficacy of fusion and control proteins, five mice per dose group received a single intraperitoneal (ip) injection with saline, darbepoetin, or Targeted EPO in a 200 μL volume (diluted in saline or PBS) on Day 0. 1-5 μL of whole blood was collected by tail-nick in EDTA-coated tubes on days 0, 4, and 7 post-injection. Blood was analyzed immediately after collection by flow cytometry as described before.[4] Thiazole orange (Sigma-Aldrich) was used to stain residual RNA in reticulocytes and reticulated platelets, and anti-CD41-PE antibody (BD Pharmingen) was used to stain total platelets. A stock solution (1 mg/mL) of thiazole orange was prepared in 100% methanol and was diluted 1:5,000 in PBS to make a 2x working solution. Anti-CD41-PE antibody was diluted 1:500 in either the 2x working solution of thiazole orange for stained samples or PBS for gating thiazole orange-negative population. 2 μL of whole blood was diluted 1:1,000 in 2 mL of PBS. Equal volumes (100 μL) of 2x working solution of anti-CD41-PE antibody with or without thiazole orange and diluted whole blood were mixed in a 96-well U-bottom plate and incubated for 30 min in the dark at RT. The fluorescence was measured on a LSRFortessa SORP flow cytometer equipped with an optional HTS sampler (BD Biosciences) using the following filter configuration: PE excitation, 561/50 mW; emission filter, BP 582/15; YFP excitation, 488/100 mW; emission filter, BP 540/25.

**Preparation of human red blood cells**

For FRAP, blood was obtained from healthy adult volunteers in accordance with the guidelines of, and approved by the Institutional Review Board of Beth Israel Deaconess Medical Center. Human blood was collected into HBSS++ containing 0.05% IgG-free BSA by venipuncture. RBCs were centrifuged at 5,000 x g at 23°C for 1 min, and were washed once with the same buffer. For flow cytometry and rosetting assays, human whole blood collected in an ACD tube by venipuncture was purchased from Zen-Bio (Research Triangle Park, NC). RBCs were isolated by centrifugation at 5,000 x g at 23°C for 1 min, and washed twice with PBS-glucose (PBS, 5 mM glucose, pH 7.4).

**Fluorescence recovery after photobleaching (FRAP)**

RBCs (10% v/v in HBSS++ containing 0.1% IgG-free BSA) were incubated with lipid dye DiO (15 μg/mL) for 30 min at 23°C in dark and washed once with HBSS++ containing 0.1% IgG-free BSA. Then RBCs were treated with GPA antibodies at 37°C for 15 min. The concentration of antibody was determined by its $K_D$ to achieve 80% saturation of GPA. RBCs were imaged using a 60x objective on an Olympus BX62 fluorescence microscope, bleached with Vector Controller Laser, and analyzed as described before.[6, 7] Reported data represent mean ± 95% confidence interval of 8-9 cells (experiment 1) or 16-20 cells (experiment 2). Data from each experiment were normalized to the buffer control in the same experiment to account for day-to-day variability, and the normalized values were combined across experiments. Mann-Whitney test was performed to determine p-values for individual and combined data sets.

**Flow cytometry for EPO accessibility and RBC crosslinking measurement**

The fluorescence was measured on a LSRFortessa SORP flow cytometer equipped with an HTS sampler (BD Biosciences) using the following filter configuration: PE excitation, 561/50 mW; emission filter, BP 582/15. Data were analyzed using FlowJo Data Analysis software (TreeStar, Ashland, OR). Reported data represent mean ± SEM of three replicates.

**RBC rosette assay**

A2780 and MCF-7 cells were plated at $2.0x10^5$ cells/mL in a 2 mL volume on a 12 x 12 mm coverslip (Electron microscopy sciences) in a well of a 6-well plate (Corning) and incubated overnight at 37°C in 5% $CO_2$. The following day, a mixture of RBCs (0.2% v/v) and fusion proteins was prepared in binding medium (RPMI-1640 without bicarbonate) and incubated for 30 min at 23°C. Coverslips coated with cells were washed twice with PBS and transferred to a new 6-well plate. The RBC-fusion protein mixture (200 μL) was added on top of each coverslip and incubated for 1 hr at 37°C. Coverslips were gently washed twice with PBS. Rosettes were imaged on a Nikon Eclipse TE300 inverted phase-contrast microscope (Nikon, Melville, NY), using a Retiga EXi CCD camera (QImaging, Surrey, Canada) controlled by iVision 4.7 software (BioVision). Rosettes were defined as adherent cells bound by 3 or more RBCs and were shown as a percentage of at least 160 cells from at least 8 fields. Reported data represent mean ± SEM of at least two replicates.

**Mouse tail vein bleeding time measurement**

The tail transection assay of bleeding time was generally performed on mice as follows. On day 0, mice were injected with a test protein (or vehicle). When testing an unknown protein, it is important to also include mice that are injected with saline or PBS vehicle as a negative control, and EPO or darbepoetin as a positive control. Typically 10 mice per dose group were used. The experiments were performed in a blind manner: one experimenter performed the injections of proteins into the mice, maintained the key, performed the injection of anaesthetic, and then handed the mice in a random order to a second, blinded experimenter. The second experimenter performed the tail transections and measured the bleeding time. For most experiments, on day 1 the tail transection and bleeding time was measured. In some experiments, the experiment was performed on day 4 so that changes in reticulocytes and reticulated platelets could also be measured. However, the advantage of performing the measurement 1 day after treatment is that after only 24 hours, the level of circulating red blood cells will not have changed, so effects on blood clotting are due to direct effects on some element of the clotting system, and not due to changes in blood viscosity.

Tail transection was performed as follows. The mice were first anaesthetized using anaesthetics ketamine and xylazine. These anaesthetics are chosen because they are thought to not affect blood clotting. Acepromazine is not used because it has the effect of reducing clotting. Mice were weighed, and mice were then injected with 120-160 mg ketamine /kg and 10-16 mg xylazine /kg of body weight. For older and heavier mice, sometimes an additional injection of about 25% of the first injection was required. After a mouse became unresponsive to a stimulus such as significant pressure to a hind foot, the mouse was placed on a heated pad on a platform over a water bath. The water bath was maintained at $37^oC$. 50-mL blue-cap tubes (Sarstedt) were filled with 50 mL of a solution of 0.85 to 0.9% NaCl that has been equilibrated to $37^oC$ in a separate water bath. The animal was placed on a Chux pad for the transection.

A position on the tail that is 3 mm from the tip, not counting hair, was marked with a felt-tip pen using calipers. (The tail was also inspected for signs of bruising that may be due to fighting, and data from such a mouse was discarded if the transected tail did not bleed at all. The decision to discard the data was always made in a blinded manner at the time of measurement). The tail was transected with a flat razor blade using a section of the blade that has not been used previously. Within 1-2 seconds, the transected tail was placed in a tube with pre-warmed saline, and then the body of the mouse was placed on the heated pad above the water bath. At the moment that the tail was transected by the blinded experimenter, the non-blinded experimenter started a timer.  The body of the mouse was then positioned on the heated pad so that only the tip of the tail – about 0.5 to 2 mm – is in the saline and the rest is in the air.  When observing the bleeding tail, the tube was rotated so that the white stripe was behind the tail, providing contrast, and the room was well-lit.  The rack holding the 50-mL tube was white or yellow to provide contrast.  The bleeding time was recorded by noting when bleeding stops, and then observing the submerged tail for up to one minute.  If bleeding re-started within this minute, the first recorded time was not counted.  Bleeding may stop and re-start several times.  If the tail is still bleeding when 10 minutes have elapsed, the time was recorded as 10 minutes.

The median and mean bleeding times were calculated for each treatment group.  Calculating the median has the advantage that extreme events, such as 10-minute time points, do not disproportionately contribute to the calculation.  Data from each experiment were normalized to the vehicle control in the same experiment to account for day-to-day variability. The normalized values were combined across experiments and Mann-Whitney test was performed to determine p-values the combined data set.

**Supplementary Tables**

Table S1. Summary of ability of diverse forms of Targeted EPO to stimulate proliferation of TF-1 cells *in vitro*. Bold indicates data shown in Figure 2b.

| Protein (V region–linker–EPO) | N | Activity in vitro Log(EC50) ± SD | EC50 relative to epoetin alpha |
|---|---|---|---|
| **Epoetin alfa (Amgen)** | **16** | **-10.239 ± 0.345** | **1** |
| **Darbepoetin (Amgen)** | **8** | **-8.989 ± 0.135** | **14.67** |
| **EPO(WT)** | **3** | **-10.378 ± 0.528** | **0.87** |
| **EPO(R150A)** | **7** | **-8.078 ± 0.233** | **130.86** |
| **EPO(K45D)** | **3** | **-6.572 ± 0.754** | **7298** |
| **10F7-35-EPO(R150A)** | **4** | **-8.947 ± 0.206** | **16.71** |
| 10F7-29-EPO(R150A) | 6 | -9.320 ± 0.303 | 7.78 |
| 10F7-18-EPO(R150A) | 5 | -9.173 ± 0.297 | 10.89 |
| 10F7-17-EPO(R150A) | 6 | -9.163 ± 0.306 | 11.27 |
| 10F7-7-EPO(R150A) | 5 | -9.386 ± 0.253 | 6.39 |
| **10F7-5-EPO(R150A)** | **8** | **-9.378 ± 0.754** | **13.69** |
| **10F7-EPO(R150A)-HEL** | **4** | **-9.142 ± 0.501** | **16.02** |
| **10F7-EPO(R150A)-LEH** | **4** | **-8.648 ± 0.643** | **54.76** |
| **1C3-35-EPO(R150A)** | **6** | **-10.517 ± 0.471** | **0.81** |
| 1C3-EPO(R150A)-HEL | 4 | -10.682 ± 0.900 | 0.98 |
| 1C3-EPO(R150A)-LEH | 4 | -10.808 ± 0.527 | 0.30 |
| **R18-17-EPO(R150A)** | **2** | **-9.415 ± 0.409** | **6.44** |
| R18-7-EPO(R150A) | 2 | -10.074 ± 0.690 | 1.95 |
| **R18-5-EPO(R150A)** | **2** | **-9.495 ± 0.185** | **4.56** |
| R18-17-EPO(K45D) | 2 | -8.828 ± 0.786 | 39.24 |
| R18-7-EPO(K45D) | 1 | -8.494 ± N/A | 43.71 |
| R18-5-EPO(K45D) | 1 | -8.019 ± N/A | 130.58 |
| **IH4-35-EPO(R150A)** | **16** | **-10.385 ± 0.756** | **1.45** |
| IH4-17-EPO(R150A) | 5 | -10.243 ± 0.505 | 1.34 |
| IH4-7-EPO(R150A | 5 | -10.479 ± 0.645 | 1.13 |
| **IH4-5-EPO(R150A)** | **9** | **-10.410 ± 0.293** | **0.69** |
| **IH4-35-EPO(K45D)** | **5** | **-9.591 ± 0.364** | **4.70** |
| IH4-5-EPO(K45D) | 1 | -9.559 ± N/A | 3.76 |

Table S2. Summary of epitopes and binding kinetics of anti-GPA antibody fragments and EPO (wildtype and R150A mutant) studied in this work.

| Protein | $k_{on}$ ($M^{-1}s^{-1}$) | $k_{off}$ ($s^{-1}$) | $K_D$ (nM) | Antibody form | GPA epitope | References |
|---|---|---|---|---|---|---|
| 10F7 | – | – | 95 | Fab | $_{34}$YAATP$_{38}$ | Chasis *et al*, 1988<br>Chasis & Mohandas, 1992<br>Catimel *et al.*, 1993 |
| 1C3 | – | – | 230 | Fab | – | Catimel *et al.*, 1993 |
| | – | – | 62 | scFv | – | Patent Application WO1994007921 |
| R18 | – | – | 25 | IgG | $_{49}$RTVY$_{52}$ | Gardner *et al.*, 1989 |
| | – | – | 400 | Fab | | |
| IH4 | $5.73 \times 10^5$ | 0.019 | 33.72 | Nanobody (VHH) | $_{52}$YPPE$_{55}$ | Habib *et al.*, 2013 |
| EPO(WT) | $3.9 \times 10^4$ | $2.1 \times 10^{-4}$ | 5.4 | N/A | N/A | Burrill *et al.*, 2016 |
| EPO(R150A) | $4.2 \times 10^4$ | $3.4 \times 10^{-3}$ | 81 | N/A | N/A | Burrill *et al.*, 2016 |

Table S3. A list of DNA sequences encoding anti-GPA antibody fragments, linkers, and EPO.

| | Sequence | Reference |
|---|---|---|
| IH4 nanobody | CAGGTCCAACTGCAGGAGAGCGGCGGGGGGTCAGTTCAGGCGGGGGGGAGTCTGCGGTTGAG<br>CTGCGTAGCTTCAGGCTACACTGACAGCACCTACTGCGTGGGATGGTTTCGGCAGGCACCCG<br>GCAAGGAACGAGAGGGCGTTGCACGGATCAACACTATCTCCGGTCGGCCTTGGTACGCAGAT<br>AGTGTTAAGGGACGGTTTACTATTAGTCAGGATAACTCTAAGAATACCGTCTACCTTCAGAT<br>GAATAGCCTGAAACCGGAAGACACGGCTATTTACTATTGCACCCTTACAACTGCCAACAGCA<br>GAGGGTTTTGTTCTGGGGGATATAACTACAAAGGACAGGGGACCCAAGTCACTGTCAGC | |
| IH4* nanobody | CAGGTCCAACTGCAAGAGAGCGGAGGAGGGTCTGTTCAAGCTGGCGGTTCCCTCCGGCTTTC<br>TTGCGTGGCGTCAGGCTATACTGACAGCACATACTGCGTGGGCTGGTTCAGGCAGGCCCCTG<br>GAAAGGAGCGCGAGGGCGTAGCCCGCATAAATACTATATCTGGCAGACCGTGGTACGCTGAC<br>AGCGTGAAGGGACGGTTTACAATCAGTCAAGATAACTCTAAAAACACCGTGTTTCTTCAAAT<br>GAATTCTTTGAAACCCGAAGATACTGCCATCTATTATTGCACACTTACGACCGCGAACTCAC<br>GCGGTTTTTGTAGCGGAGGATATAACTATAAAGGGCAAGGGCAGGTAACTGTATCC | |
| 10F7 scFv | CAAGTTAAGTTGCAACAATCTGGTGCTGAATTGGTTAAGCCAGGTGCTTCTGTTAAGTTGTC<br>TTGTAAGGCTTCTGGTTACACCTTCAACTCTTACTTTATGCATTGGATGAAGCAAAGACCAG<br>TTCAAGGTTTGGAATGGATTGGTATGATTAGACCAAACGGTGGTACTACCGATTACAACGAG<br>AAGTTTAAGAACAAGGCTACTTTGACTGTTGATAAGTCCTCTAACACTGCTTACATGCAATT<br>GAACTCTTTGACTTCTGGTGATTCTGCTGTTTACTACTGTGCTAGATGGGAAGGTTCTTACT<br>ACGCTTTGGATTACTGGGGTCAAGGTACCACTGTTACTGTTTCTTCCGGTGGAGGTGGATCT<br>GGTGGTGGAGGATCTTCAGGAGGTGGTGGATCTTCCGATATTGAGTTGACTCAATCTCCAGC<br>TATTATGTCTGCTACCTTGGGTGAGAAGGTTACTATGACTTGTAGAGCTTCATCTAACGTTA<br>AGTACATGTACTGGTACCAACAGAAGTCTGGTGCTTCTCCAAAGTTGTGGATTTACTACACT<br>TCTAACTTGGCTTCTGGTGTTCCAGGTAGATTTTCTGGTTCAGGTTCTGGTACTTCCTACTC<br>TTTGACTATTTCCTCTGTTGAAGCTGAAGATGCTGCTACTTACTACTGTCAACAATTCACTT<br>CTTCCCCATACACTTTTGGAGGAGGTACTAAGTTGGAAATCAAG | GenBank (accession no. KX026660-3) |
| 1C3 scFv | GAAGTCCGTCTGCTGGAAAGCGGGGGTGGTCCTGTGCAGCCTGGTGGGTCCCTGAAACTGTC<br>CTGTGCCGCAAGCGGGTTCGATTTTTCCAGATACTGGATGAACTGGGTGAGGAGGGCTCCAG<br>GCAAGGGCCTGGAGTGGATCGGCGAGATCAACCAGCAGTCCAGCACCATCAATTACTCTCCC<br>CCTCTGAAGGACAAGTTCATCATCAGCCGCGATAACGCTAAGTCTACACTGTATCTGCAGAT<br>GAATAAGGTGAGAAGCGAGGACACCGCCCTGTACTATTGCGCTCGCCTGTCTCTGACAGCCG<br>CTGGCTTTGCCTATTGGGGCCAGGGCACCCTGGTGACAGTGTCTGCTGGAGGAGGCTCTTCC<br>GGAGGATCCGGCAGCTCTGGCGGCTCCAGCTCTGGCGGCGATATCGTGATGAGCCAGTCTCC<br>CTCCAGCCTGGCCGTGTCCGTGGGAGAGAAGGTGTCCATGAGCTGTAAGTCTTCCCAGTCTC<br>TGTTCAACTCCAGAACCCGCAAGAATTACCTGACATGGTATCAGCAGAAGCCTGGCCAGAGC<br>CCCAAGCCTCTGATCTACTGGGCCAGCACCAGAGAGTCTGGAGTGCCAGACCGCTTCACCGG<br>CTCTGGATCCGGCACAGACTTCACCCTGACAATCAGCTCTGTGCAGGCCGAGGACCTGGCTG<br>ATTACTATTGCAAGCAGTCCTATAATCTGAGGACCTTTGGCGGCGGCACAAAGCTGGAGATC<br>AAG | |
| R18 scFv | CAGGTTAAACTCCAGCAAAGTGGTGGCGGGCTCGTACAACCAGGCGGTTCCCTCAAGTTGTC<br>CTGCGCCGCATCAGGGTTTACATTTAGCTCTTATGGTATGTCTTGGTTTCGCCAGACGCCTG<br>ACAAGCGACTCGAGCTGGTCGCTATCATCAATAGTAACGGAGGTACTACATATTATCCCGAC<br>AGTGTGAAGGGGCGATTTACCATTAGCCGGGACAACGCCAAAAATACACTGTACCTCCAGAT<br>GTCAAGCTTGAAATCAGAAGATACGGCCATGTACTATTGCGCTAGGGGGGGTGGAAGGTGGC<br>TTCTGGACTATTATGGTCAGGGTACAACAGTGACAGTATCCTCCGGTGGAGGTGGATCTGGT<br>GGTGGAGGATCTTCAGGAGGTGGTGGATCTTCCGACATAGAGCTTACACAATCTCCGTCATC<br>ACTGGCAGTCTCAGCCGGGGAAAAAGTGACAATGTCATGCAAGTCAAGCCAGAGCGTTCTTT<br>ATTCATCTAATCAGAAGAACTACCTGGCATGGTATCAGCAGAAGCCGGGACAGTCCCCTAAG<br>CTCCTCATCTACTGGGCAAGCACCAGGGAATCCGGAGTGCCGGACAGGTTTACTGGGTCCGG<br>TTCTGGGACGGATTTTACGCTTACGATATCAAGTGTCCAAGCTGAGGACCTCGCAGTATACT<br>ACTGTCACCAGTACCTGTCTTCTTCTACTTTTGGGGGTGGAACGAAACTGGAAATAAAA | |
| EPO (WT) | GCTCCACCTAGATTGATTTGTGATTCCAGAGTTTTGGAAAGATACTTGTTGGAAGCTAAGGA<br>GGCTGAAAATATTACTACTGGTTGTGCTGAACATTGTTCTTTGAACGAGAATATTACTGTTC<br>CAGATACTAAGGTTAACTTTTACGCTTGGAAGAGAATGGAAGTTGGTCAGCAAGCTGTTGAA<br>GTTTGGCAAGGTTTGGCTTTGTTGTCTGAAGCTGTTTTGAGAGGTCAAGCTTTGTTGGTTAA<br>TTCTTCTCAACCATGGGAACCATTGCAATTGCATGTTGATAAGGCTGTTTCTGGTTTGAGAT<br>CTTTGACTACCTTGTTGAGAGCTTTGGGTGCTCAAAAGGAAGCTATTTCTCCTCCAGATGCT<br>GCTTCTGCCGCTCCATTGAGAACTATTACTGCTGATACTTTTAGAAAGTTGTTTAGAGTTTA<br>CTCTAACTTCTTGAGAGGTAAGTTGAAGTTGTACACTGGTGAAGCTTGTAGAACTGGTGATC<br>GG | GenBank (accession no. KX026660-3) |
| EPO (R150A) | GCTCCACCTAGATTGATTTGTGATTCCAGAGTTTTGGAAAGATACTTGTTGGAAGCTAAGGA<br>GGCTGAAAATATTACTACTGGTTGTGCTGAACATTGTTCTTTGAACGAGAATATTACTGTTC | GenBank (accession |

| | | |
|---|---|---|
| | CAGATACTAAGGTTAACTTTTACGCTTGGAAGAGAATGGAAGTTGGTCAGCAAGCTGTTGAA<br>GTTTGGCAAGGTTTGGCTTTGTTGTCTGAAGCTGTTTTGAGAGGTCAAGCTTTGTTGGTTAA<br>TTCTTCTCAACCATGGGAACCATTGCAATTGCATGTTGATAAGGCTGTTTCTGGTTTGAGAT<br>CTTTGACTACCTTGTTGAGAGCTTTGGGTGCTCAAAAGGAAGCTATTTCTCCTCCAGATGCT<br>GCTTCTGCCGCTCCATTGAGAACTATTACTGCTGATACTTTTAGAAAGTTGTTTAGAGTTTA<br>CTCTAACTTCTTGGCCGGTAAGTTGAAGTTGTACACTGGTGAAGCTTGTAGAACTGGTGATC<br>GG | no.<br>KX026660-3) |
| EPO<br>(K45D) | GCTCCACCTAGATTGATTTGTGATTCCAGAGTTTTGGAAAGATACTTGTTGGAAGCTAAGGA<br>GGCTGAAAATATTACTACTGGTTGTGCTGAACATTGTTCTTTGAACGAGAATATTACTGTTC<br>CAGATACTGATGTTAACTTTTACGCTTGGAAGAGAATGGAAGTTGGTCAGCAAGCTGTTGAA<br>GTTTGGCAAGGTTTGGCTTTGTTGTCTGAAGCTGTTTTGAGAGGTCAAGCTTTGTTGGTTAA<br>TTCTTCTCAACCATGGGAACCATTGCAATTGCATGTTGATAAGGCTGTTTCTGGTTTGAGAT<br>CTTTGACTACCTTGTTGAGAGCTTTGGGTGCTCAAAAGGAAGCTATTTCTCCTCCAGATGCT<br>GCTTCTGCCGCTCCATTGAGAACTATTACTGCTGATACTTTTAGAAAGTTGTTTAGAGTTTA<br>CTCTAACTTCTTGAGAGGTAAGTTGAAGTTGTACACTGGTGAAGCTTGTAGAACTGGTGATC<br>GG | GenBank<br>(accession<br>no.<br>KX026660-3) |
| 5AA | TCTGGTGGTGGTTCC | |
| 7AA | GGAGGATCTGGTGGTGGTTCC | |
| 17AA | GGAGGATCCGGTGGTGGAGGATCATCTGGTGGAGGATCTGGTGGTGGTTCC | |
| 29AA | GGAGGAAGTTCCGGTGGTGGATCTTCTTCTGGAGGTGGAGGATCCGGTGGTGGAGGATCATC<br>TGGTGGAGGATCTGGTGGTGGTTCC | |
| 35AA | GGTGGAGGTGGTTCCGGAGGAGGAAGTTCCGGTGGTGGATCTTCTTCTGGAGGTGGAGGATC<br>CGGTGGTGGAGGATCATCTGGTGGAGGATCTGGTGGTGGTTCC | |

Table S4. A list of protein sequences for anti-GPA antibody fragments, linkers, and EPO.

| | Sequence | Reference |
|---|---|---|
| IH4 nanobody | QVQLQESGGGSVQAGGSLRLSCVASGYTDSTYCVGWFRQAPGKEREGVARINTISGRPWYAD<br>SVKGRFTISQDNSKNTVYLQMNSLKPEDTAIYYCTLTTANSRGFCSGGYNYKGQGTQVTVS | US patent 9879090[1] |
| IH4* nanobody | QVQLQESGGGSVQAGGSLRLSCVASGYTDSTYCVGWFRQAPGKEREGVARINTISGRPWYAD<br>SVKGRFTISQDNSKNTVFLQMNSLKPEDTAIYYCTLTTANSRGFCSGGYNYKGQGQVTVS | US patent 9879090[1] |
| 10F7 scFv | QVKLQQSGAELVKPGASVKLSCKASGYTFNSYFMHWMKQRPVQGLEWIGMIRPNGGTTDYNE<br>KFKNKATLTVDKSSNTAYMQLNSLTSGDSAVYYCARWEGSYYALDYWGQGTTVTVSSGGGGS<br>GGGGSSGGGGGSSDIELTQSPAIMSATLGEKVTMTCRASSNVKYMYWYQQKSGASPKLWIYYT<br>SNLASGVPGRFSGSGSGTSYSLTISSVEAEDAATYYCQQFTSSPYTFGGGTKLEIK | |
| 1C3 scFv | EVRLLESGGGPVQPGGSLKLSCAASGFDFSRYWMNWVRRAPGKGLEWIGEINQQSSTINYSP<br>PLKDKFIISRDNAKSTLYLQMNKVRSEDTALYYCARLSLTAAGFAYWGQGTLVTVSAGGGSS<br>GGSGSSGGSSSGGDIVMSQSPSSLAVSVGEKVSMSCKSSQSLFNSRTRKNYLTWYQQKPGQS<br>PKPLIYWASTRESGVPDRFTGSGSGTDFTLTISSVQAEDLADYYCKQSYNLRTFGGGTKLEI<br>K | Patent application WO1993024630[2] |
| R18 scFv | QVKLQQSGGGLVQPGGSLKLSCAASGFTFSSYGMSWFRQTPDKRLELVAIINSNGGTTYYPD<br>SVKGRFTISRDNAKNTLYLQMSSLKSEDTAMYYCARGGGRWLLDYYGQGTTVTVSSGGGGSG<br>GGGSSGGGGGSSDIELTQSPSSLAVSAGEKVTMSCKSSQSVLYSSNQKNYLAWYQQKPGQSPK<br>LLIYWASTRESGVPDRFTGSGSGTDFTLTISSVQAEDLAVYYCHQYLSSSTFGGGTKLEIK | US patent 8900592[8] |
| EPO (WT) | APPRLICDSRVLERYLLEAKEAENITTGCAEHCSLNENITVPDTKVNFYAWKRMEVGQQAVE<br>VWQGLALLSEAVLRGQALLVNSSQPWEPLQLHVDKAVSGLRSLTTLLRALGAQKEAISPPDA<br>ASAAPLRTITADTFRKLFRVYSNFLRGKLKLYTGEACRTGDR | |
| EPO (R150A) | APPRLICDSRVLERYLLEAKEAENITTGCAEHCSLNENITVPDTKVNFYAWKRMEVGQQAVE<br>VWQGLALLSEAVLRGQALLVNSSQPWEPLQLHVDKAVSGLRSLTTLLRALGAQKEAISPPDA<br>ASAAPLRTITADTFRKLFRVYSNFLAGKLKLYTGEACRTGDR | |
| EPO (K45D) | APPRLICDSRVLERYLLEAKEAENITTGCAEHCSLNENITVPDTDVNFYAWKRMEVGQQAVE<br>VWQGLALLSEAVLRGQALLVNSSQPWEPLQLHVDKAVSGLRSLTTLLRALGAQKEAISPPDA<br>ASAAPLRTITADTFRKLFRVYSNFLRGKLKLYTGEACRTGDR | |
| 5AA | SGGGS | |
| 7AA | GGSGGGS | |
| 17AA | GGSGGGGSSGGGSGGGS | |
| 29AA | GGSSGGGSSSGGGGSGGGGSSGGGSGGGS | |
| 35AA | GGGGSGGGSSGGGSSSGGGGSGGGGSSGGGSGGGS | |

**Supplementary Figures**

a

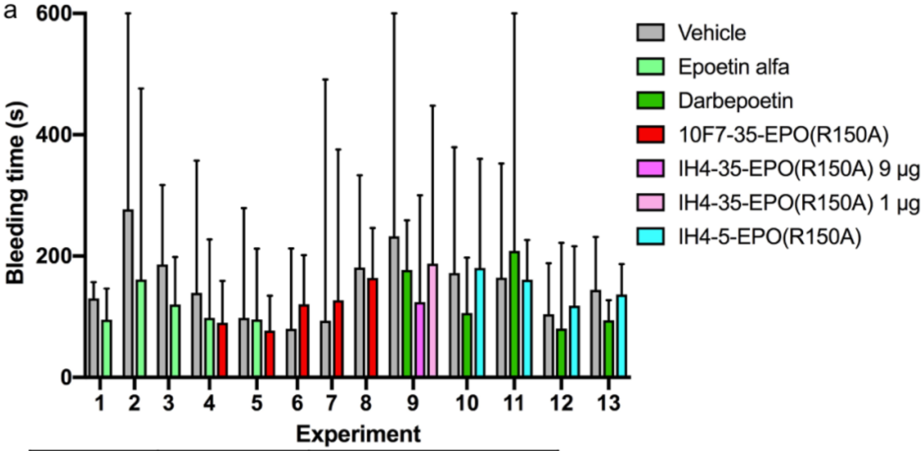


b

| Experiment | Human GPA | Dose groups |
|---|---|---|
| 1<br>2<br>3 | Non-transgenic | Vehicle<br>Epoetin alfa |
| 4<br>5 | Transgenic | Vehicle<br>Epoetin alfa<br>10F7-35-EPO(R150A) |
| 6<br>7<br>8 | Non-transgenic | Vehicle<br>10F7-35-EPO(R150A) |
| **9* | *Transgenic* | *Vehicle*<br>*Darbepoetin*<br>*IH4-35-EPO(R150A) 9 µg*<br>*IH4-35-EPO(R150A) 1 µg* |
| 10<br>11<br>12<br>13 | Transgenic | Vehicle<br>Darbepoetin<br>IH4-5-EPO(R150A) |

c

| Protein (V region–linker–EPO) | Dose (pmol) | Human GPA | N | Median bleeding time relative to vehicle ± SD | p-value | Significance |
|---|---|---|---|---|---|---|
| Epoetin alfa (Amgen) | 50 | Transgenic | 2 | 0.840 ± 0.190 | 0.0063 | *** |
| | 50 | Non-transgenic | 3 | 0.653 ± 0.074 | | |
| Darbepoetin (Amgen) | 50 | Transgenic | 4 | 0.828 ± 0.301 | 0.0207 | ** |
| 10F7-35-EPO(R150A) | 125 | Transgenic | 2 | 0.717 ± 0.098 | 0.0527 | * |
| | 125 | Non-transgenic | 3 | 1.257 ± 0.313 | 0.1750 | ns |
| IH4-5-EPO(R150A) | 40 | Transgenic | 4 | 1.028 ± 0.083 | 0.8015 | ns |
| **Darbepoetin (Amgen)* | *50* | *Transgenic* | *1* | *0.761 ± N/A* | *0.2855* | *ns* |
| **IH4-35-EPO(R150A)* | *180* | *Transgenic* | *1* | *0.533 ± N/A* | *0.1200* | *ns* |
| | *20* | *Transgenic* | *1* | *0.806 ± N/A* | *0.7815* | *ns* |

Figure S1. Summary of bleeding times after treating huGPA-transgenic or non-transgenic mice with various forms of non-targeted and targeted EPO. Mice received a single ip injection of darbepoetin or epoetin alfa (50 pmol; 1.8 µg), 10F7-EPO(R150A) (125 pmol; 9 µg), IH4-5-EPO(R150A) (40 pmol; 2 µg), or vehicle on day 0, and bleeding times were measured on day 1. In experiment 9 (italics, *), mice received two ip injections of darbepoetin (50 pmol; 1.8 µg), IH4-35-EPO(R150A) (180 pmol; 9 µg or 20 pmol; 1 µg), or vehicle on days 0 and 3, and bleeding times were measured on day 4. (a) Raw data from all 13 bleeding time experiments. Data represent median with interquartile range. (b) Description of each experiment shown in (a). (c) Bleeding times were normalized to the median of a vehicle control performed on the same day, to account for day-to-day variability. The normalized values were combined across experiments and Mann-Whitney test was performed. *p < 0.1; **p < 0.05; ***p < 0.01; ns: not significant.

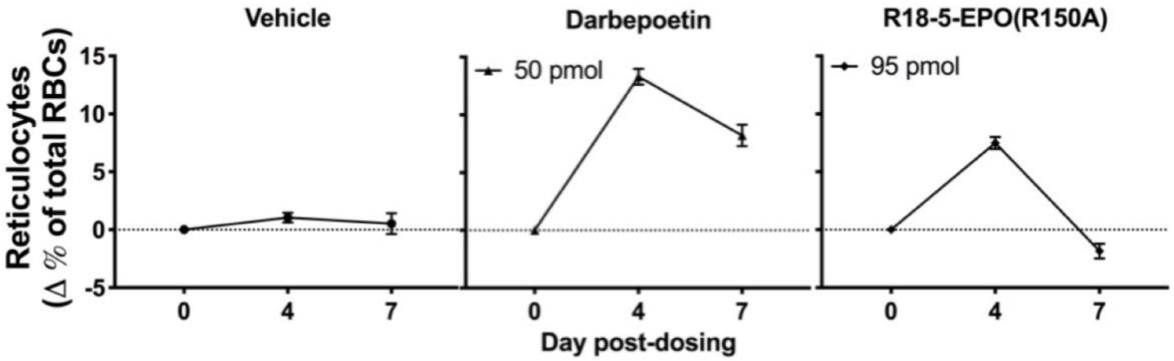


Figure S2. Ability of a form of Targeted EPO, R18-5-EPO(R150A), to promote RBC production in human GPA-transgenic mice. Compared to IH4-5-EPO(R150A), a higher dose of R18-5-EPO(R150A) is needed to induce comparable increase in reticulocytes.

a

```
                      <---CDR1--->                <-------CDR2------->
      1         11         21         31         41         51         61
      .......... .......... .......... .......... .......... .......... ...
IH4   QVQLQESGGG SVQAGGSLRL SCVASGYTDS TYCVGWFRQA PGKEREGVAR INTISGRPW  YADSVKGRFT
IH4*  QVQLQESGGG SVQAGGSLRL SCVASGYTDS TYCVGWFRQA PGKEREGVAR INTISGRPW  YADSVKGRFT
1G6V  QVQLVESGGG SVQAGGSLRL SCAASGYTVS TYCMGWFRQA PGKEREGVAT ILGGSTY    YGDSVKGRFT
4W6X  QVQLQESGGG SVQAGGSLRL SCTASGYTYR KYCMGWFRQA PGKEREGVAC INSGGGTSY  YADSVKGRFT
5HDO  QVQLQESGGG LVQAGGSLRL SCAASGFTLD SYAIGWFRQA PGKEREGVSC ISASGGSTN  YADSVKGRFT
5MY6  QVQLQESGGG SVQAGGSLKL TCAASGYIFN SCGMGWYRQS PGRERELVSR ISGDGDTWHK   ESVKGRFT
2X6M   GQLVESGGG SVQAGGSLRL SCAASGIDSS SYCMGWFRQR PGKEREGVAR INGLGGVKTA YADSVKDRFT
1OP9  QVQLQESGGG SVQAGGSLRL SCSASGYTY     ISGWFRQA PGKEREGVAA IRSSDGTTY  YADSVKGRFT

                                 <---------CDR3---------->
      71        81         91         101        111        121        131
      .......... .......... .......... .......... .......... .......... ...
IH4   ISQDNSKNTV YLQMNSLKPE DTAIYYCTLT T  ANSRGFC SG      GYN YKGQGTQVTV S
IH4*  ISQDNSKNTV FLQMNSLKPE DTAIYYCTLT T  ANSRGFC SG      GYN YKGQG QVTV S
1G6V  ISQDNAKNTV YLQMNSLKPE DTAIYYCAGS T  VASTGWC SR LRPYDYH YRGQGTQVTV SS
4W6X  ISQDNAKDTV FLRMNSLKPE DTAIYYCALS SNSVCPPGHV A      WYN DWGQGTQVTV SS
5HDO  ISRDNAKNTV YLQMNSLKSE DTAVYYCAAD H  PGLCTSE SGR RRYLE  VWGQGTQVTV SSA
5MY6  ISQDNVKKTL YLQMNSLKPE DTAVYFCA        VCYN        LET YWGQGTQVTV SS
2X6M  ISRDNAENTV YLQMNSLKPE DTAIYYCAAK FS PG  Y C GGSWSNFGY   WGQGTQVTV SSH
1OP9  ISQDNAKNTV YLQMNSLKPE DTAMYYCAAT E  VA  GW    PLDIGIYD YWGQGTEVTV SS
```

b

1G6V: Dromedary nanobody to bovine carbonic anhydrase
4W6X: Llama nanobody to lectin domain of F18 fimbrial adhesin FedF
5HDO: Dromedary nanobody to urokinase-type plasminogen activator
5MY6: Dromedary nanobody to HER2
2X6M: Dromedary nanobody to alpha-synuclein
1OP9: Camelid nanobody to human lysozyme

Figure S3. Protein sequence alignments of IH4 and other nanobodies. (a) The original IH4 sequence (IH4*) was modified to match the consensus sequence. IH4* (blue) indicates the original protein sequence of the IH4 nanobody from the US Patent No. 9,879,090.[1] Phe81 in the framework region 3 of IH4* is mutated to tyrosine (green highlight), and a threonine residue is inserted between Gly117 and Gln118 in the framework region 4 of IH4* (green highlight). The resulting sequence is shown as IH4 (orange, bold). Each dot indicates a position for a single amino acid residue. Numbers indicate amino acid positions every 10 residues. Complementarity-determining regions (CDRs) are shown with dots highlighted in gray and underlined sequences. Sequences between CDRs are framework regions. (b) PDB ID's and brief descriptions of six nanobodies used in sequence alignments.

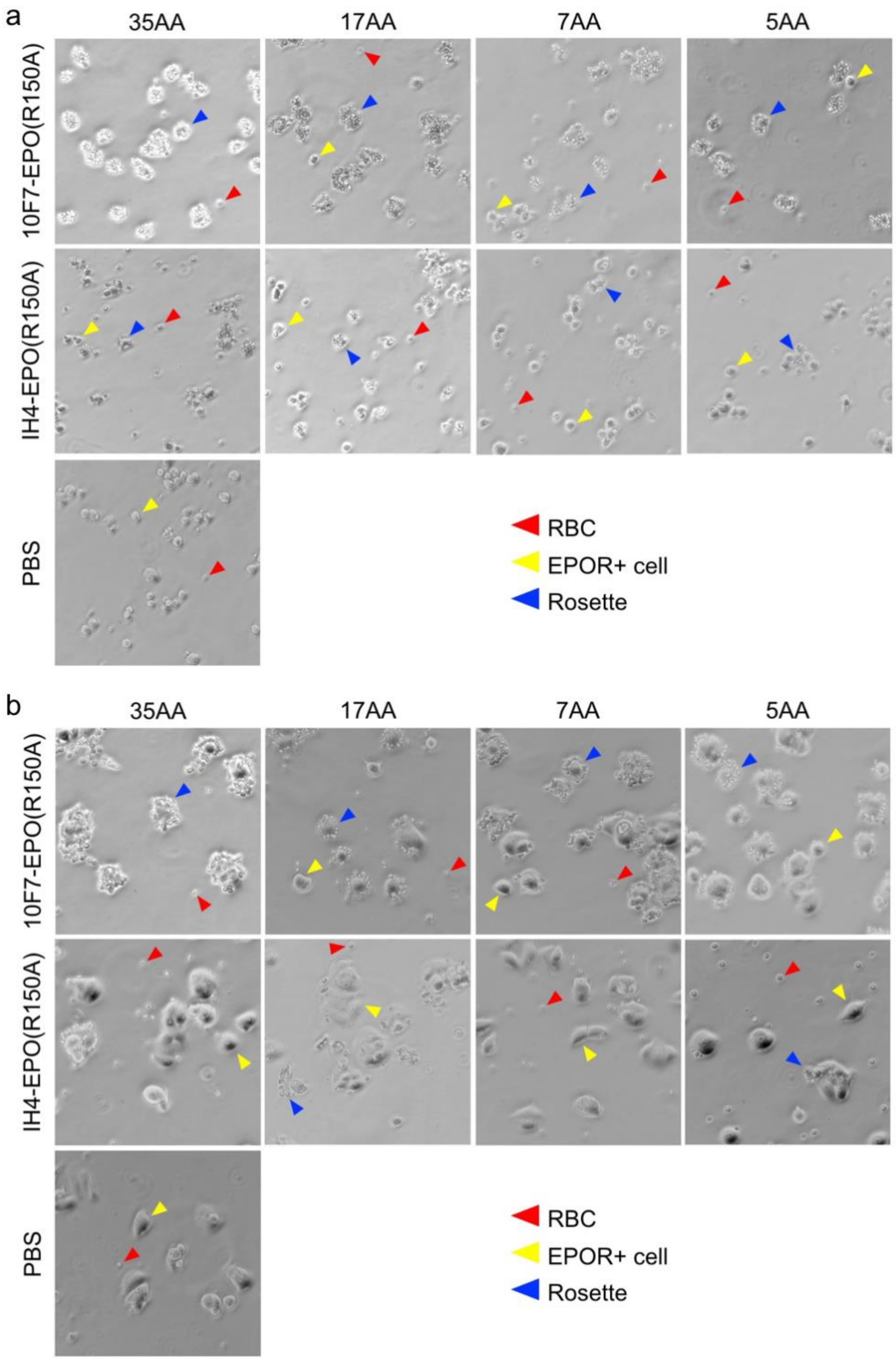


Figure S4. Phase-contrast images of RBC rosetting around EPO-R-bearing cells. Human RBCs treated with Targeted EPO variants were incubated with (a) A2780 or (b) MCF-7 cells. 200x magnification.

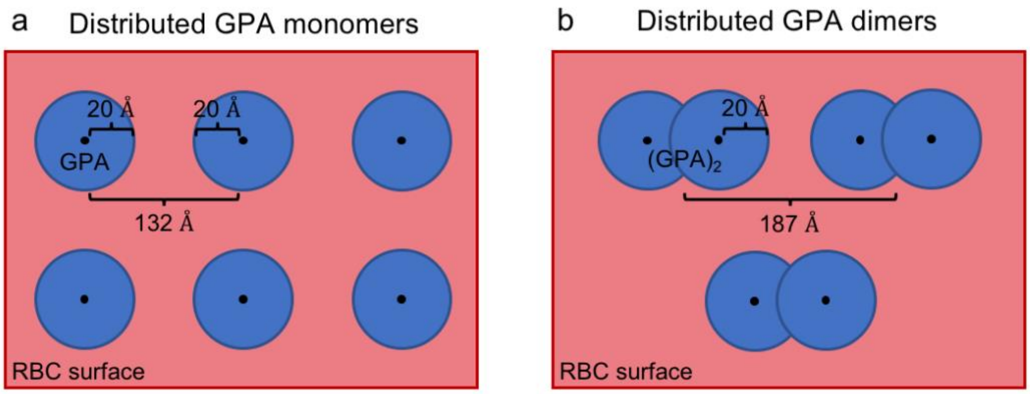


c

| | Equation | GPA monomers | GPA dimers |
|---|---|---|---|
| RBC surface area | – | 140 µm$^2$ | |
| The number of GPA molecules | – | 800,000 | 400,000 |
| The number of GPA molecules per µm$^2$ | $\frac{The\ number\ of\ GPA\ molecules}{RBC\ surface\ area}$ | 5714.3 | 2657.2 |
| Distance (Å) between GPA molecules | $\sqrt{(The\ number\ of\ GPA\ molecules\ per\ \mu m^2)^{-1}}$ | 132.3 | 187.1 |

Figure S5. Geometric model of the RBC surface. There are 800,000 GPA monomers densely packed on the RBC surface, whose area is about 140 µm$^2$. The average distance between GPA molecules is about 130-190 Angstroms (Å). (a) When all GPA molecules are monomeric, the average distance between the two adjacent monomers is about 132 Å. (b) When all GPA molecules dimerize, the average distance between the two adjacent dimers is about 187 Å. O-linked glycans extend 20 Å from the peptide backbone. (c) Summary of relevant numbers and equations to estimate the density and accessibility of GPA molecules on the RBC surface.
